# Delineation of a novel assembly intermediate in retroviral integration pathway

**DOI:** 10.1101/2025.09.09.675266

**Authors:** Rahul Chadda, Sibes Bera, Mohamed Ghoneim, Tamara De Melo, Duane P. Grandgenett, Edwin Antony, Krishan K. Pandey

## Abstract

Retroviral integration is mediated by viral integrase (IN) which synapses two viral long terminal repeat (LTR) DNA ends and produces a series of nucleoprotein complexes known as intasomes. While structural studies of mature intasomes have illuminated key aspects of their architecture and provided insights into the integration reaction, the sequence of events driving IN oligomerization and engagement of the viral DNA pairing remains unclear. Here, using complementary biochemical and biophysical approaches, including ensemble and single molecule FRET, we reveal that integration progresses through a key transient intermediate that leads to the mature intasome. We demonstrate that Rous Sarcoma Virus (RSV) intasome assembly pathway proceeds through a tetrameric intermediate where two IN dimers engage a single DNA end. This complex subsequently oligomerizes to form mature functional octameric intasome in which two DNA ends are juxtaposed for concerted integration. These findings provide mechanistic insights into the stepwise pathway of retroviral integration and define a previously uncharacterized intermediate critical for intasome maturation.

**Graphical abstract:** 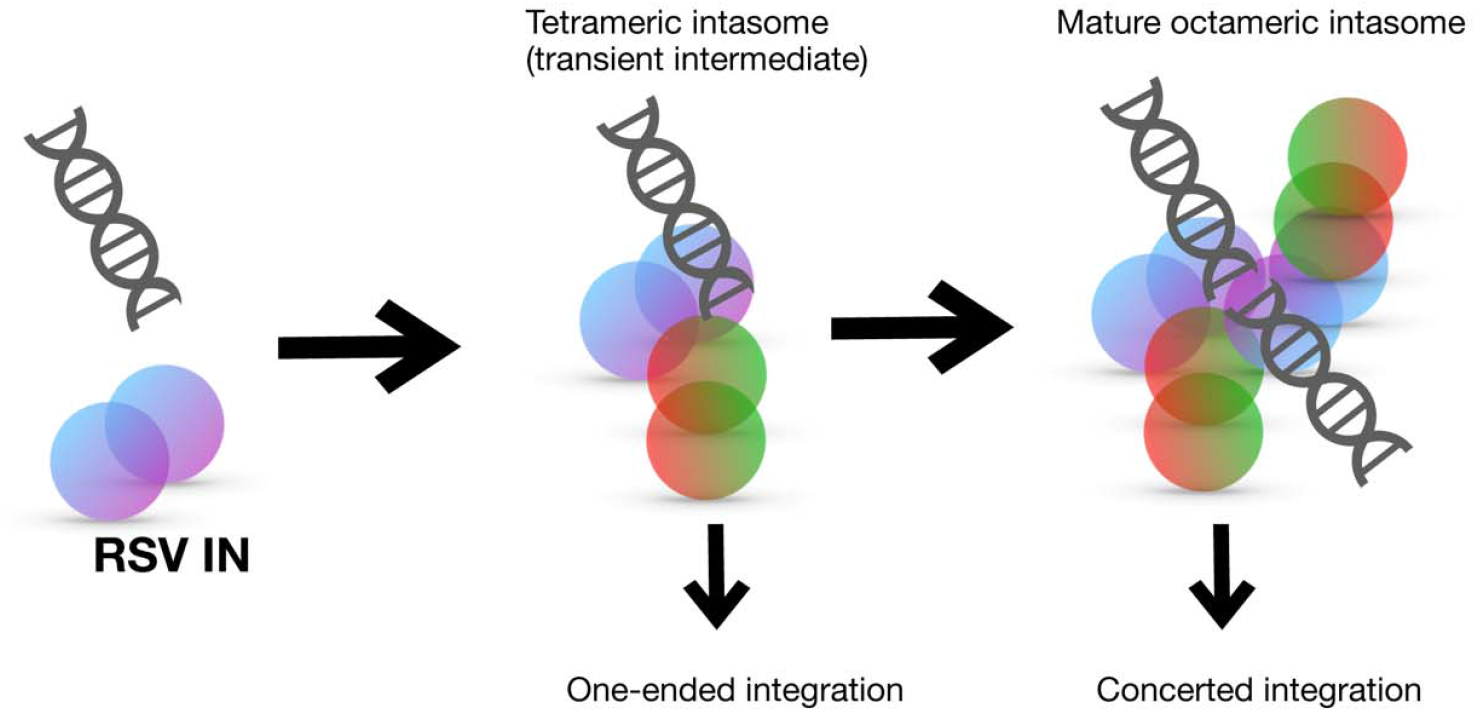

## Introduction

Retrovirus integrase (IN) possesses a unique ability to insert the ends of linear viral DNA in a concerted fashion into the host genome, a step required for virus replication. Despite advances in our understanding of integration from structural studies, there is a substantial gap in deciphering the assembly mechanisms associated with these IN/viral DNA complexes, here termed intasomes. IN from Rous sarcoma virus (RSV), HIV-1, and related retroviruses share a conserved three-domain architecture composed of a catalytic core domain (CCD) that is flanked by amino (NTD) and carboxy-terminal domains (CTD) (1-3). The NTD adopts a three-helix bundle and contains the conserved HHCC motif that coordinates a Zn^2+^ ion (4, 5). The CCD is the most conserved domain among retroviruses and contains the active site comprising the DDE (Asp-Asp-Glu) motif which is responsible for catalytic activity (6, 7). The CTD, which adopts an SH3-like fold is the least conserved domain (8). The CTD is proposed to be the primary factor driving formation of different oligomeric forms of nucleoprotein complexes observed with different retroviral INs (9, 10). The structure of the flexible C-terminal tail region which extends beyond the CTD has not been fully resolved. The 17 aa tail region (270-286 aa) in RSV IN has a key role in defining intasome architecture (11-13).

During integration, IN binds both ends of linear viral DNA (generated by reverse-transcription) and removes two nucleotides from the 3’-termini of catalytic strand producing cleaved synaptic complex (CSC). IN within the CSC intasome binds host DNA to form the strand transfer complex (STC) and mediates the nicking of host DNA and joining of viral DNA to host DNA. In general, most intasome assemblies observed in vitro suggest DNA-mediated tetramerization of the predominant IN species observed in solution although the precise oligomeric states vary across retroviruses. Intasomes are composed of IN multimers ranging from 4 subunits for prototype foamy virus (PFV) (14), human T-cell leukemia virus (HTLV-1) (15) and simian T-lymphotropic virus type 1 (16) to 8 for alpha-retrovirus Rous sarcoma virus (RSV) (17, 18) and beta-retrovirus mouse mammary tumor virus (MMTV) (19, 20). Lentiviral intasomes display greater variability, containing between 4 and 16 IN subunits (21-24). Thus, intasomes exhibit considerable structural diversity despite conserved catalytic function.

Available intasome structures represent end-products of the integration pathway; the CSC formed with IN and viral LTR DNA and the STC formed with a branched DNA substrate mimicking viral LTR DNA covalently joined to the target DNA. We previously showed that the C-terminal tail region of RSV IN plays a critical role in determining the CSC intasome architecture (**Fig. 1, 2**). RSV IN (1-269), lacking the tail region, predominantly forms the tetrameric intasome (**Fig. 1B, 2**) whereas full-length RSV IN (1-286) forms the octameric intasome (11). IN (1-278) was the most efficient in producing octameric intasomes as end-products (12) (**Fig 1A, 2**). Interestingly, truncations within the C-terminal tail region (270-286 aa) did not significantly affect the 3’-processing (12), strand transfer activities and their ability to produce STC in presence of a target DNA **(Supp Fig. S1, Fig 1C)**. Strand transfer and 3’-processing activity is typically measured at 37^°^C while the intasome assembly is carried out at 18^°^C to prevent catalysis.

**Fig. 1.**
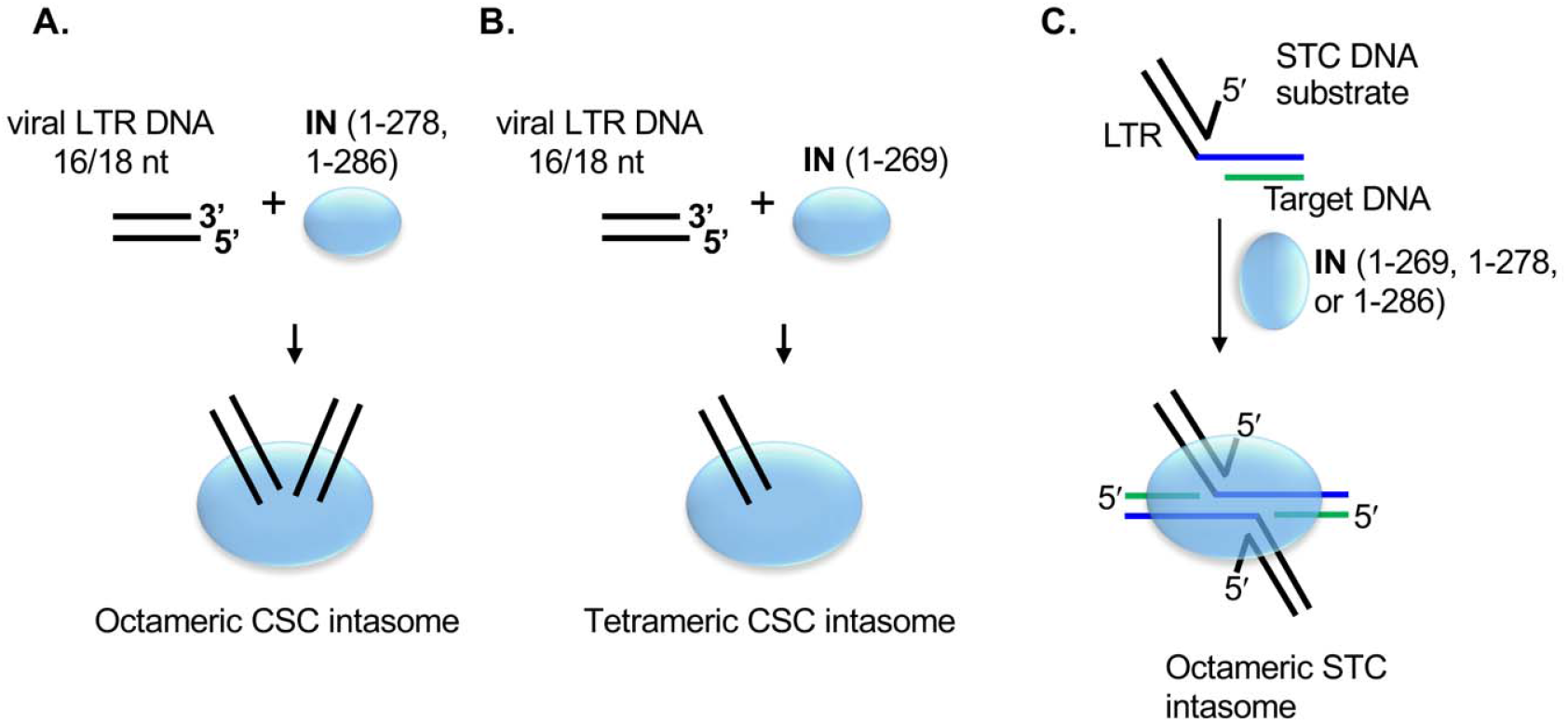
RSV intasome assembly showing the terminal IN-DNA complexes produced in vitro. **A**. The wt full-length RSV IN (1-286 aa) or the IN (1-278) having an 8 aa truncation in the C-terminal tail region binds to viral LTR DNA ends, resulting in the assembly of a stable octameric CSC. **B**. IN (1-269) having a truncation of complete 17 aa tail region and certain missense IN mutants produce a tetrameric CSC intasome only. **C**. Strand transfer complex (STC) can be assembled in vitro using a branched STC substrate mimicking viral DNA covalently joined to target DNA. All of RSV INs produce octameric STC intasome. Multimeric forms of IN are not shown in these schematics.

**Fig. 2.**
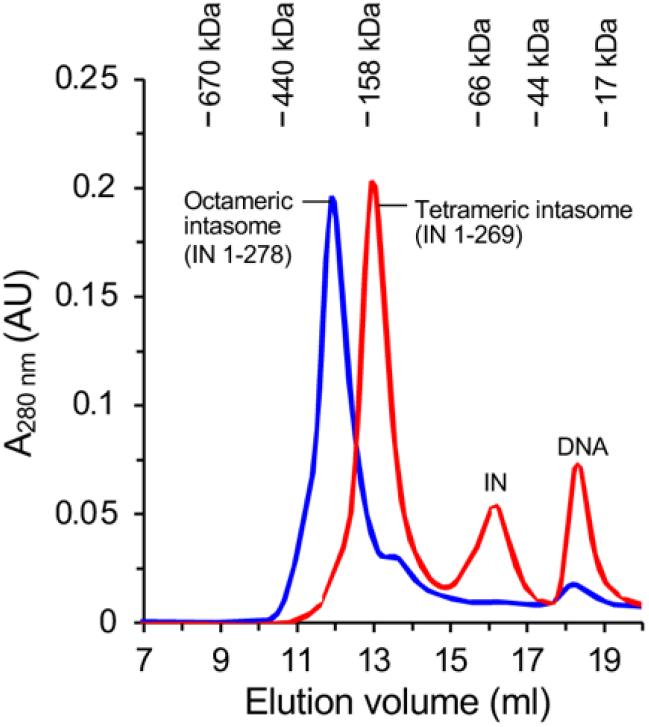
Differential Octameric assembly of RSV intasomes. and tetrameric intasomes are exclusively end-products formed with IN (1-278 aa) or IN (1-269 aa).

Our preliminary studies indicated that mature octameric intasome formation with IN (1-278) and IN (1-286) was preceded by a tetrameric intermediate (12). We hypothesize that there are key intermediates in the intasome assembly pathway with distinct IN multimers and viral DNA that subsequently results in the binding of target DNA and integration. We proposed that RSV tetrameric intasome may be a transient precursor to the mature octameric intasome. In this study, we further establish the steps associated with formation of tetrameric intasome precursor and its conversion to mature octameric intasome. Our results show that the precursor tetrameric intasome consists of two IN dimers bound to a single DNA molecule which subsequently oligomerizes to yield the functional mature octameric intasome. These findings provide mechanistic insight into the stepwise assembly of retroviral intasomes and identify a previously uncharacterized intermediate that is critical for integration.

## Materials and Methods

### RSV IN expression and purification

RSV IN constructs were expressed in *Escherichia coli* BL21 (DE3) pLysS and purified to near-homogeneity as described previously (11, 13, 25). The wild-type (wt) Prague A IN subunit is 286 aa in length, designated IN (1-286). Purified IN was concentrated to 20-30 mg/ml using Amicon Ultra-15 centrifugal filters (30K MWCO). For this study, RSV IN (1-278 aa) is simply referred to as RSV IN and remains predominantly dimeric in solution (**Supp Fig. S2**). The protein concentrations are reported as monomeric subunits. The DNA sequence of all IN constructs was confirmed by sequencing.

### Concerted integration assay

The concerted integration assays were performed using 3’-OH recessed oligonucleotide viral DNA substrates and RSV IN as described previously (11, 12). Double-stranded 3’-OH recessed substrates containing RSV gain-of-function(G) U3 were synthesized by Integrated DNA Technologies. The DNA substrates were recessed by two nucleotides on the catalytic strand and designated with “R”. The oligonucleotide length denotes the non-catalytic strand. The sequences were as follows: 18R GU3 (5’-ATT GCA TAA GAC **<u>A</u>**AC A-3’ and 5’-AAT GT**<u>T</u>** GTC TTA TGC AAT-3’). The bold underlined nucleotide on the catalytic strand differed between the GU3 and wt U3 sequence. The concentrations of IN and the viral oligonucleotides in a typical assay were 2 and 1 µM, respectively. The strand transfer products were resolved on a 1.3% agarose gels, stained with SYBR Gold (Invitrogen) and analyzed using a Typhoon 9500 Laser Scanner (GE Healthcare Life Sciences).

### Assembly of the RSV cleaved synaptic complex (CSC)

RSV CSC intasomes were assembled with IN and GU3 18R in the presence or absence of integrase strand transfer inhibitor (INSTI) MK-2048 (11). The intasomes were analyzed by size-exclusion chromatography (SEC) using Superdex 200 Increase (10/300) at 4°C. The intasome assembly without INSTI was performed in 20 mM HEPES, pH 7.5, 150 mM NaCl, 50 mM MgSO_4_, 1 M non-detergent sulfobetaine (NDSB)-201, 10% dimethyl sulfoxide (DMSO), 10% glycerol, and 1 mM tris(2-carboxyethyl phosphine)(TCEP). Intasomes were purified by SEC in 20 mM HEPES pH 7.5, 150 mM NaCl, 50 mM MgSO_4_, 5% glycerol and 1 mM TCEP. The assembly reactions which contained INSTI MK-2048 were carried out in 20 mM HEPES, pH 7.5, 100 mM NaCl, 100 mM ammonium sulfate, 1 M non-detergent sulfobetaine (NDSB)-201, 10% dimethyl sulfoxide (DMSO), 10% glycerol, 1 mM TCEP. INSTI bound intasomes were purified by SEC in 20 mM HEPES pH 7.5, 200 mM NaCl, 100 mM ammonium sulfate, 5% glycerol and 1 mM TCEP. Reaction mixture typically contained 45 μM IN (as monomers), and 15 μM 3’ OH recessed oligonucleotide and where indicated 125 μM MK-2048. MK-2048 was generously provided by Merck & Co. The samples were incubated at 18°C for 18 h unless otherwise indicated. For time-course experiments, the half-time (t_1/2_) of the tetramer to octamer conversion was determined by non-linear fit using equation “Y=Y_0_ + (Plateau-Y_0_) (1-exp^-Kx^)”; where Y_0_ is the initial value, plateau is the Y value at 24 h, K is the rate constant, expressed in reciprocal of X unit (time). The half-time (t_1/2_) was computed as ln(2)/K.

### Mass-photometry

The SEC purified fractions containing tetrameric and octameric fractions were analyzed by a TwoMP instrument (Refeyn Inc) as described (26). Briefly, the purified complexes were diluted 1:10 in 20 mM HEPES, 100 mM NaCl, 200 mM ammonium sulfate, 1 mM TCEP pH 7.5 and immediately applied on a clean slide for light scattering measurements. Molecular weight standards were resuspended in matching buffer and used to calibrate and convert the particle-image contrast due to scattered light into MW units. Standards included β-amylase (3 species of 56, 112, & 224□kDa, respectively) and thyroglobulin (single species of 667□kDa).

### Steady-state Fluorescence (bulk-FRET)

RSV IN intasomes were assembled as described earlier using viral LTR DNA internally labeled at position 9 with Cy3 (iCy3-18R GU3; 5’-AAT GTT GTC /iCy3/TTA TGC AAT-3’) or Cy5 and (iCy5-18R GU3; 5’-AAT GTT GTC /iCy5/TTA TGC AAT-3’) on the non-transferred strand and annealed with complementary transferred strand (5’-ATT GCA TAA GAC AAC A-3’). Equimolar quantities (7.5 μM each) of iCy3 18R-GU3 (donor)- and iCy5 18R-GU3 (acceptor)-labeled annealed 18R GU3 were used in the assembly mixture. Labeling at position 9 was not expected to affect IN binding or intasome assembly as determined from the IN-DNA interactions in cryo-EM structures (18, 27). The distance between 9^th^ nucleotide on two DNA ends within the CSC intasome is 39 Å which is appropriate for F□rster resonance energy transfer (FRET) analysis (18, 27). The complexes were assembled under conditions suitable to allow purification of tetrameric as well as octameric intasomes from the same sample. Intasome assembly and concerted integration was not affected by internal fluorophore labeling of viral LTR DNA. Octameric and tetrameric intasomes, purified by SEC, were analyzed using a Fluoromax-3 (Jobin Yvon, Inc., Edison, NJ) spectrofluorometer at 14 °C with a temperature-regulated cell holder. The samples were excited with 550 nm (donor excitation), and emission spectra were collected from 555 to 720 nm for quenching of Cy3 (peak at□565 nm) and sensitized emission of Cy5 (peak at□668 nm). All spectra were corrected for buffer and instrument.

### Single-molecule FRET (smFRET)

The SEC purified fractions containing tetrameric or octameric fractions were analyzed by smFRET-TIRFm (total internal reflection fluorescence microscopy). The complexes were assembled with equimolar ratio of iCy3 18R-GU3 (donor)- and iCy5 18R-GU3 (acceptor)-labeled 18R GU3 (7.5 μM each) with RSV IN (45 μM). The tetrameric and octameric intasomes were purified using SEC and immediately analyzed by smFRET. Intasome complexes were loaded onto pre-cleaned coverslip-chambers. The complexes were allowed to settle down passively. Excess molecules were washed off using binding buffer (20 mM HEPES pH 7.5, 200 mM NaCl, 100 mM ammonium sulfate and 1 mM TCEP). Single molecule TIRF imaging was performed in the same buffer supplemented with oxygen scavenging system and Trolox using an inverted, objective based total-internal-reflection fluorescence microscope (TIR-FM; IX71 Olympus). Samples were excited with a 532nm laser or 637nm laser to excite the donor or the acceptor, respectively. Donor fluorescence (green traces) and energy-transferred acceptor traces (red) were collected at the rate of 100 ms frame time using an EM-CCD (iXon Ultra DU-897U-CS0). The data analysis was carried out as described earlier (28).

To analyze the freely diffusing intasomes in solution, smFRET data were collected on an EI-FLEX bench-top microscope (Exciting Instruments Ltd Sheffield, UK) as described (29) with minor modifications. The intasomes were assembled using equimolar iCy3 and iCy5 18R GU3 with or without INSTI MK-2048 and purified by SEC as above. Before measurement on EI-FLEX, respective SEC buffers were photo irradiated using 100W LED bulb for 3 days in cold room (4°C). Buffers were supplemented with 1 mM TCEP and 0.1 mg/ml photo irradiated BSA for intasome dilutions just before the measurement. Octamer and tetramer intasome were diluted to 10 pM on ice. A 100 μl sample droplet was placed onto a no.1 thickness coverslip and excited with alternating 520 nm (0.22 mW) and 638 nm (0.15 mW) lasers respectively. Lasers were sequentially turned ON for 45 μs for each measurement and separated by a dark period of 5 μs for a total of 30 mins of acquisition at room temperature (21°C). Fluorescence emission photons from freely diffusing molecules were collected using an Olympus 60× (1.2 N.A.) water-immersion objective, focused onto a 20 μm pinhole. After passing through the pinhole, the photons were split using a 640 nm long-pass filter, cleaned up using 572 and 680 nm band-pass filters, and focused onto respective avalanche photodiodes. The photon arrival times, and respective detector were saved in HDF5 data format for offline analysis. Several measurements were recorded with freshly diluted samples for each intasome populations. The data was analyzed in Anaconda environment (2.6.6), with Jupyter notebooks, using FRETBursts Python package 0.7.1 (30). FRET values of single molecule bursts of photon emissions were exported in Excel sheets then plotted as FRET histograms in GraphPad Prism 10.

### Stopped-flow analysis of IN-DNA interactions

To monitor the initial interactions between IN and DNA leading to the assembly of CSC, we performed stop flow experiments to monitor the IN-DNA (iCy3-18R GU3) dynamics using an Applied Photophysics SX20 (UK) instrument in reaction buffer containing 20□mM HEPES, pH 7.5, 125□mM NaCl, 10□mM MgCl_2_, 10% DMSO (v/v) and 5 mM DTT. The change in photoisomerization-induced fluorescence enhancement (PIFE) of Cy3 was measured upon binding to IN. 100 nM iCy3-18RGU3 DNA and varying amount of RSV IN (20 nM – 300 nM) were rapidly mixed and monitored by exciting the samples at 530□nm, and emission was measured using a 555 nm cut-off filter. Protein and DNA concentrations denoted here are pre-mixing conditions, which are reduced to half after mixing the DNA and IN to provide final post-mixing concentrations. The protein induced fluorescence enhancement was monitored as IN binding enhanced the Cy3 fluorescence signal and fitted using a single exponential equation and the rate of change was plotted as a function of IN concentration (**Supp Fig. S3**). The association constant (k_ON_) and dissociation rate constant (k_OFF_) were determined to be 5.6 x 10^8^ M^-1^s^-1^ and 8.86 s^-1^, respectively by fitting the data to a linear equation. As expected, IN binds tightly to viral DNA with a dissociation constant of ~15.8 nM.

## Results

### Characterization of a novel assembly intermediate in the RSV integration pathway

Intasome structures from several retroviruses have revealed the architecture of their end-products in the concerted integration pathway. However, the pathways leading to formation of mature intasomes from IN and DNA are not well understood. Therefore, in this study, we investigated the assembly pathway using RSV IN and viral DNA, in the absence/presence of INSTI MK-2048, as a function of time (**Fig. 3, 4**). MK-2048, like other INSTIs is a strand transfer inhibitor which prevents the target DNA binding to IN-viral DNA complexes thus preventing integration (31, 32). IN remains predominantly dimeric in absence of viral DNA (**Supp Fig. S2**).

**Fig. 3.**
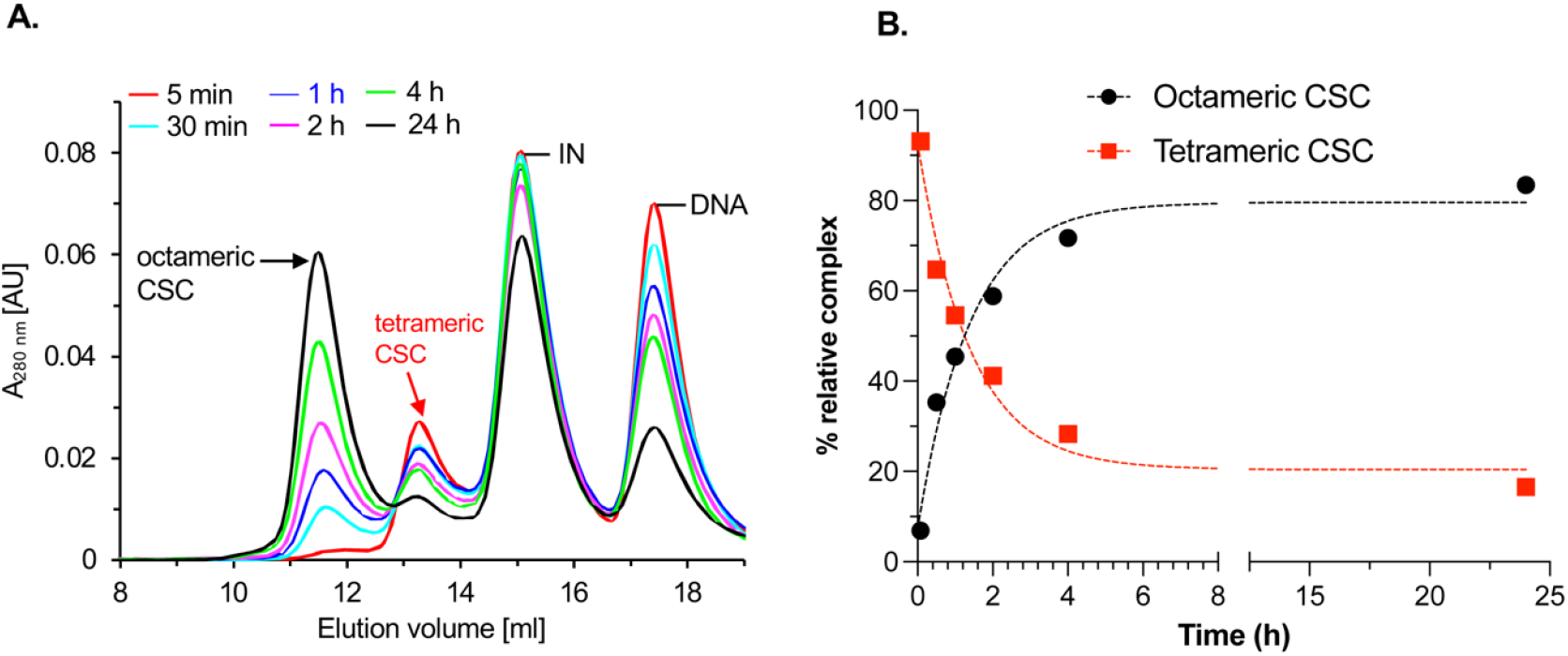
Transient RSV tetrameric CSC intasome is rapidly converted to octameric CSC intasome in absence of INSTI. **A**. SEC profiles of a time course of CSC intasome assembly with RSV IN (1-278) without INSTI. Tetrameric CSC intasomes are formed early and rapidly converted to octameric CSC intasomes. **B**. Quantitation of relative percentage of intasomes. Dotted lines indicate non-linear fit (t_1/2_ = 0.96 h).

**Fig. 4.**
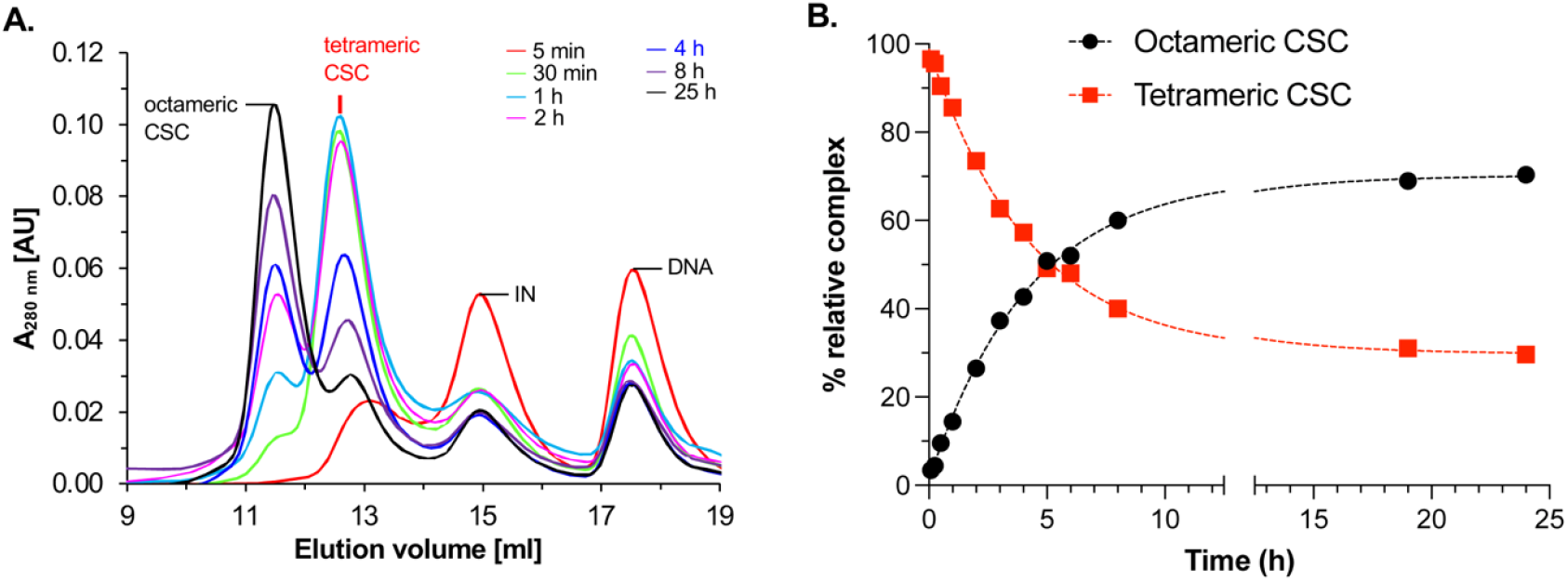
RSV tetrameric CSC intasome is an assembly intermediate for the mature octameric CSC intasome in vitro. **A**. SEC profiles for a time course of CSC intasome assembly with RSV IN (1-278) in presence of INSTI MK-2048 at 18°C. Selected assembly time points are shown for clarity. Tetrameric CSC intasomes are formed early in the assembly pathway that are gradually converted to octameric CSC intasomes. **B**. Relative percentage yield of intasome is plotted at all assembly time points. Dotted lines indicate non-linear fit (t_1/2_ = 2.94 h).

In absence of INSTI, a tetrameric intasome is formed first and gradually converted into mature octameric intasomes (**Fig. 3A, B**). The t_1/2_ for the tetramer to octamer transition is 0.96 h. In the presence of MK-2048 during assembly, the tetrameric intasome accumulates rapidly with peak yield between 30 min and 1 h and subsequently decreases with time with concomitant increase in octameric intasome assembly (**Fig. 4A, B**). The t_1/2_ for the tetramer to octamer transition is 2.94 h. Notably, INSTI was not included during intasome purification by SEC irrespective of presence or absence of INSTI during the intasome assembly. These data confirm earlier reports using HIV-1 IN (32) and RSV IN (11) that INSTI is not required for the assembly of intasomes. Rather, INSTI such as MK-2048 traps and stabilizes the intasomes in solution resulting in their higher yield.

### Pathway choice for oligomerization routes through a single DNA-bound tetrameric intasome intermediate

In the RSV integration pathway, DNA binding drives formation of the tetrameric intasome which then matures into the octameric form. Cryo-EM studies of the octameric RSV intasome revealed two DNA ends bound to opposing IN dimers (18). We have established that a DNA-bound tetrameric intasome is a precursor to the mature octamer. A structure of the tetrameric intasome precursor is not available and the mechanistic details of how it is assembled in the presence of DNA remains poorly understood. To parse this mechanism further, we started out with two possible pathways for the assembly of the RSV intasome (**Fig. 5**).

**Fig. 5.**
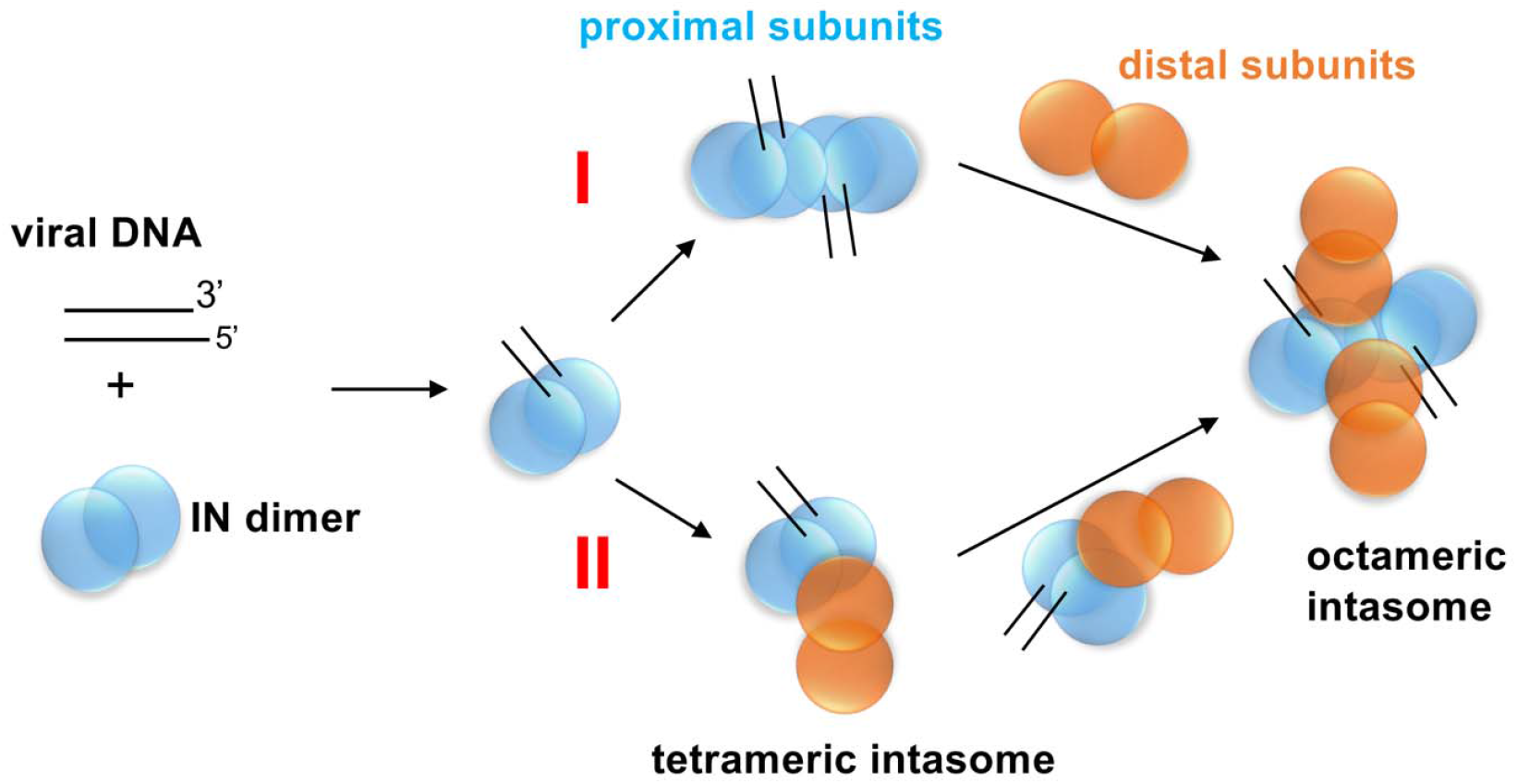
Potential assembly pathways of RSV intasome. Proximal and distal IN protomers are shown in blue and orange, respectively.

In the first pathway, two DNA -bound IN dimers (that form the proximal subunits in the final octamer) oligomerize to form an IN tetramer complex (**Fig. 5**). In this scenario, both DNA fragments are sequestered, and the subsequent steps involve engagement of the distal IN dimers leading to formation of the octameric intasome. The second pathway posits an intermediate where one DNA-bound IN dimer oligomerizes with a distal IN molecule generating a tetramer containing a single DNA which then rearranges to produce mature octameric intasome (**Fig. 5**). The key difference in the second pathway is that the two DNA fragments encounter each other only within the context of the octamer. We observed that DNA binding to RSV IN is rapid in stopped flow experiments (**Fig. S3**). In these experiments, a Cy3 labeled DNA was used and the change in Cy3 fluorescence upon binding to IN was monitored. The rate of DNA binding is much faster than the oligomerization observed in time course experiments (**Fig. 3, 4**). Therefore, with respect to pathway choice, initial IN-DNA interactions occur rapidly.

To experimentally distinguish between two pathway choices towards formation of the terminal octameric intasome, we used FRET to test the binding of two fluorescently labeled LTR DNAs with either Cy3 or Cy5. Equal quantities of both labeled DNAs were mixed with IN to assemble the intasomes and both the tetrameric and octameric intasome were separated by SEC (**Fig. 6A**). Assembly of two DNA fragments within the fractions containing tetramer and/or the octamer intasomes should give rise to a high FRET signal.

**Fig. 6.**
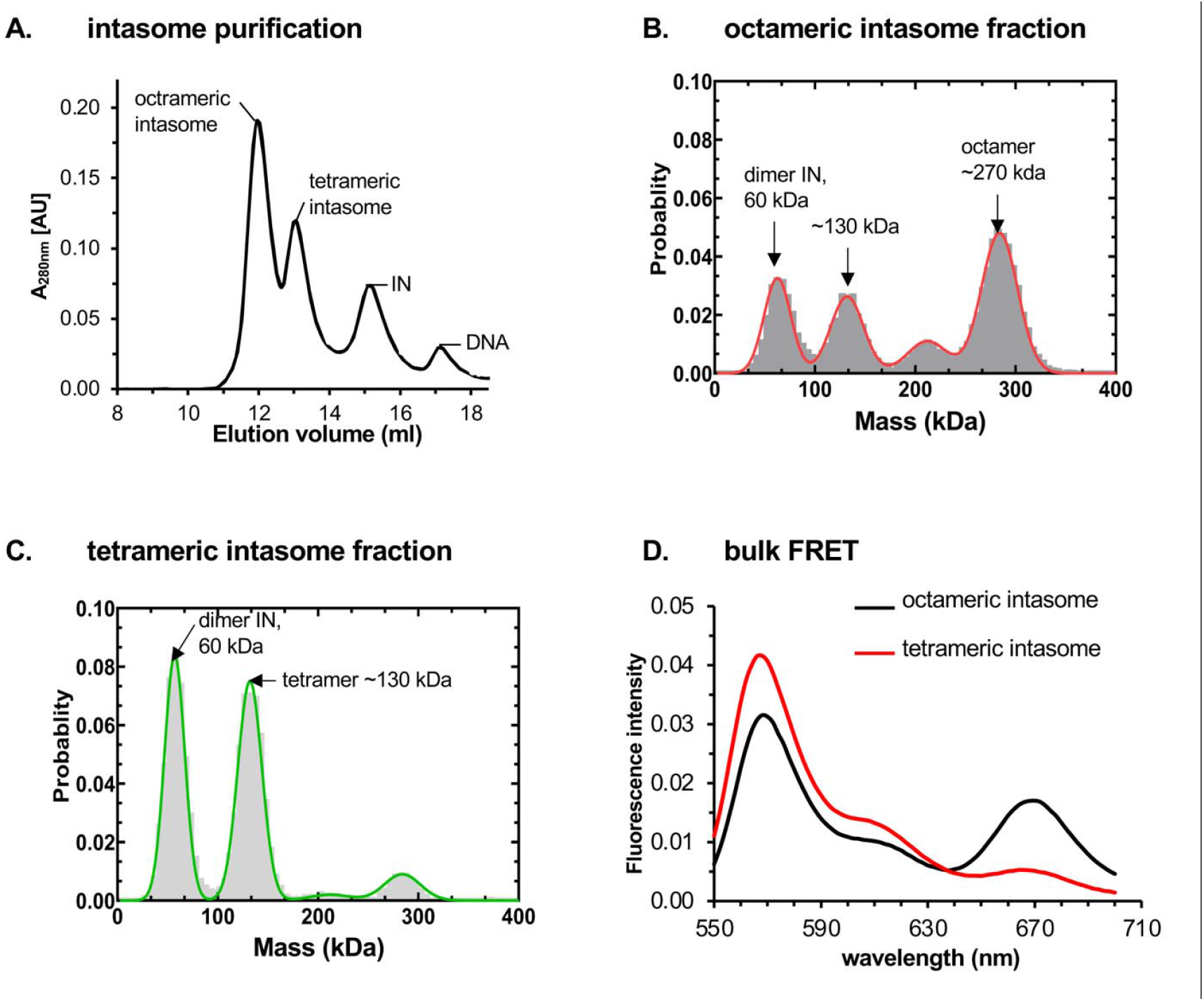
Characterization of transient tetrameric and mature octameric intasomes assembled with RSV IN. **A**. The SEC purification profile of CSC intasomes assembled with IN 1-278 and Cy3/Cy5 labeled 18R GU3 LTR DNA in presence of INSTI MK-2048. The top fractions of octameric and tetrameric intasome were analyzed by mass-photometer (B and C, respectively) and bulk FRET (D) **B. & C**. Octameric and tetrameric fractions showed molar mass predominantly of ~270 and ~130 kDa, respectively. Dimeric IN (~60 kDa) was also observed probably due to complex dissociation during analysis. **D**. Bulk-FRET of octameric and tetrameric intasomes. Normalized FRET profiles show that FRET was observed predominantly in octameric intasomes only.

The efficiency of separation of the two species was further confirmed using mass photometry. The octameric intasome fraction displayed a predominant ~270 kDa peak that corresponds to the octamer along with smaller ~60 kDa and ~130 kDa peaks that correspond to the dimeric and tetrameric species, respectively (**Fig. 6B, C**). The tetrameric intasome fraction displayed two major species that correspond to the ~60 kDa dimer and ~130 kDa tetramer, respectively, along with a very minor species for the octamer.

These tetrameric and octameric intasome fractions carrying the labeled DNA were subsequently analyzed for changes in Cy5 fluorescence signal by exciting the Cy3 donor. Fluorescence emission scanning of both samples after excitation of the Cy3 shows a robust increase in the Cy5 signal for the octameric complex compared to the tetrameric complex (**Fig. 6D**). This suggests that two DNA molecules in the octameric complex are bound and aligned as observed in the cryoEM structure (18) and positioned to generate robust FRET. In contrast, the lack of a FRET-induced Cy5 signal for the tetrameric sample suggests that this complex is composed of one DNA-bound IN dimer and a dimer of the proximal subunits not bound to DNA (pathway II; **Fig. 5**).

### Single molecule FRET confirms the one-DNA-bound tetrameric intasome intermediate

The ensemble FRET studies suggests that the transient tetrameric IN-DNA complex intermediate is likely engaged to only one DNA molecule. However, the lack of FRET-induced increase in Cy5 fluorescence could also be explained by the engagement of two DNA molecules in the tetramer but bound in an orientation that is not conducive for FRET. To rule out this possibility, we used single-molecule FRET to directly visualize the complexes and individually quantitate the co-localization of the Cy3 and Cy5 fluorescence signals. The tetrameric and octameric intasomes that were formed with Cy3/Cy5-labeled DNA in presence of INSTI MK-2048 and separated by SEC fractionation were passively coated onto cover slips and imaged using smFRET-TIRFm. We observed dynamic smFRET in only the octameric intasome fractions showing distinct transitions from high to low FRET states (**Fig. 7A**). The majority of intasome particles containing Cy3 and Cy5 labeled DNA showed FRET (**Fig. 7B**). In contrast, the majority of tetrameric intasome particles had either Cy3 or Cy5 labeled DNA individually and hence no FRET was observed (**Fig. 7B**). These results point to pathway **II** being the most plausible pathway in RSV intasome assembly. These are the first of kind single molecule studies of retroviral intasome assembly systems that characterize the intermediate IN-DNA complexes.

**Fig. 7.**
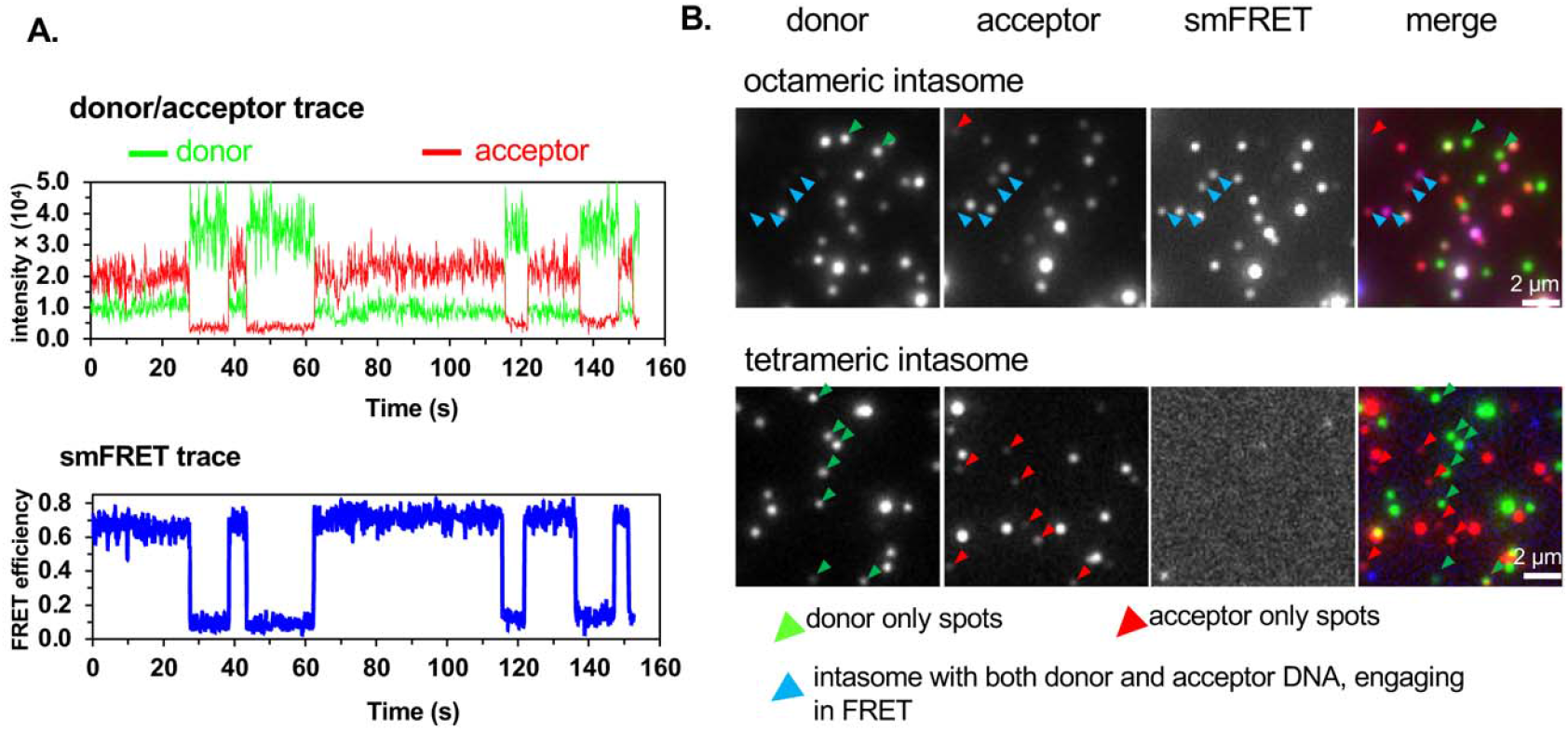
Visualization of intasome complexes by smFRET. **A**. Traces of smFRET of the octameric intasome showing dynamic FRET; the fluorescence intensity from donor (Cy3) is shown in green and acceptor (Cy5) in red while the FRET signal is shown in blue. ON/OFF smFRET due to release/recapture of DNA molecules. **B**. Snapshot of the single intasome particles in TIRF. The donor only and acceptor only spots are shown in green and red, respectively. The spots labeled with blue arrow contain both the donor and acceptor and show FRET. Please note that not all such particles are labeled.

To further quantitate these DNA binding events, we performed smFRET measurements using an EI-FLEX single molecule fluorescence spectrometer which utilizes alternating-laser excitation (ALEX) (33) to specifically sort molecules containing both donor and acceptor fluorophores. SEC purified octameric and tetrameric intasomes were diluted to 10 pM and FRET measured in-solution. The SEC purified intasomes remain stable during the measurement as evident by their re-chromatography (**Supp Fig. S4**). Lower yield of tetrameric intasome (in absence of INSTI) and its tendency to rapidly convert to mature octameric intasome did not allow us to determine its stability.

The octameric intasome had significantly higher counts showing high FRET efficiency (**Fig. 8A**) compared to the tetrameric intasome (**Fig. 8B**). These data confirm that octameric intasome has two DNA ends sequestered while tetrameric complex has just one DNA end, thus making pathway II (**Fig. 5**) as the predominant route for intasome assembly. Similar results were obtained when the intasomes were assembled in presence of INSTI MK-2048 (**Fig. 8C, D**). Tetrameric intasome formed with RSV IN (1-269) did not display FRET in a similar experiment (data not shown). It reinforces the conclusion that INSTI do not affect the intasome assembly, rather trap the intasomes in an inactive state thus preventing target DNA binding (32, 34, 35).

**Fig. 8.**
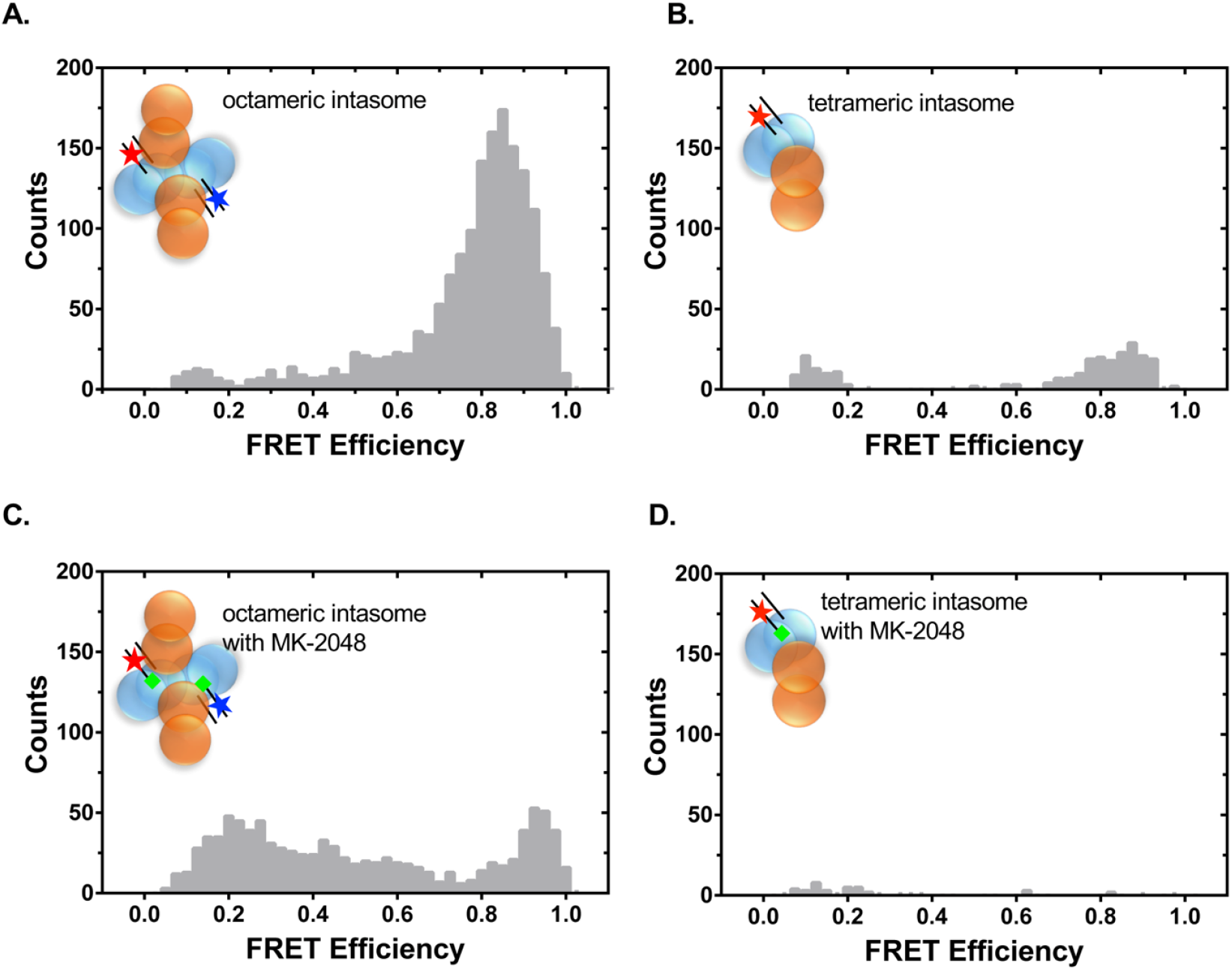
Confocal sm-FRET analysis of RSV intasomes using EI-FLEX. SEC purified intasomes were analyzed (10 pM) by EI-FLEX and number of counts showing FRET was plotted. The intasomes were assembled using equimolar concentration of iCy3 and iCy5 labeled 18RGU3 DNA. The fluorophores are indicated in red and blue. **A**. Octameric intasome without drug. **B**. Tetrameric intasome without drug. **C**. Octameric intasome formed in the presence of INSTI MK-2048. **D**. Tetrameric intasome in the presence of INSTI MK-2048. INSTI MK-2048 is shown in green. The FRET was observed predominantly in octameric intasomes only.

## Discussion

Understanding the assembly mechanisms of retrovirus intasomes by IN multimers is necessary to elucidate the pathways for viral DNA integration. The structure of different retrovirus intasomes has revealed remarkable architectural diversities besides conserved structural features and catalytic mechanisms. Generally, most intasome assemblies observed in vitro suggest DNA-mediated tetramerization of the predominant IN species in solution. It has been difficult to obtain homogeneous intasome populations with HIV-1 or HIV-2 IN primarily due to their poor solubility and tendency to form polymeric intasome complexes instead of homogeneous uniform species of complexes produced by RSV IN under our optimized solution conditions (**Fig. 2**). Here, we provide evidence that assembly of the mature RSV octameric intasome in vitro occurs via a precursor tetrameric intermediate (**Fig. 1**). The ability of RSV IN to assemble the catalytically active octameric intasome from its tetrameric precursor is unique among studied retroviral systems. Whether a transient intermediate en route to mature HIV-1 intasome exists is unknown and difficult to study due to heterogeneous intasome assembly (22, 24).

We determined the structures of the RSV octameric intasome (four IN dimers and two viral DNA molecules) (18) and strand transfer complex formed with a branched DNA substrate (27) by single particle cryo-EM. These structures as well as other retroviral intasome structures show that the two DNA ends are synapsed by opposing IN proximal subunits providing the catalytic actives sites and with their NTD ends swapped across the DNA binding interface. The tetrameric intasome by virtue of being a transient precursor to mature octameric intasome has been refractory to structural studies by cryo-EM. Hence, in this study, we utilized various in-solution approaches to define the RSV intasome assembly pathway (**Fig. 5**).

Our data support the pathway II to be the predominant route for assembly of mature octameric intasome. A FRET signal is only observed for the octameric intasomes suggesting that two DNA molecules are bound with their fluorophore labeled ends juxtaposed in proximity. The smFRET data further confirm that transient tetrameric intasomes contain only one DNA molecules. Therefore, no FRET signal is observed. Our proposed model suggests that two of these tetrameric complexes undergo rearrangement to produce mature a octameric intasome. A significant rearrangement is needed to synapse two tetrameric complexes in a pathway leading to formation of mature octameric intasomes as it requires multiple cross subunits interactions between two proximal IN subunits and two DNA ends (18, 27). Adding INSTI MK-2048 during the intasome assembly enabled us to monitor the progress of mature octameric intasome assembly from its precursor. Formation of tetramer seem to be faster than its conversion to mature octameric intasome. Within 30 minutes, the tetramer yield was maximum while its conversion to octamer was much slower process with gradual increase in its yield up to ~18 h (**Fig. 4**). INSTI allows the accumulation of trapped intasomes. INSTI binding to the tetrameric intasome slowed its conversion to mature octameric intasome as evident from t_1/2_ of 2.94 h compared to 0.96 h in absence of INSTI. INSTI binding to intasome delays and inhibit the formation of STC (3, 34). In the absence of INSTI the conversion of tetrameric to octameric intasome is faster with t_1/2_ of 0.96 h; however, the yield was lower presumably due to their lower stability.

The tetrameric complex in pathway II is ideally suited to produce circular half-site products in vitro in which only one end of viral DNA integrates into the target DNA. RSV IN (49-286) which lacks the NTD and hence unable to provide DNA binding interface to synapse opposing IN-DNA complex to produce CSC. However, it is still able to carry out single-ended integration event producing circular half site products (36). This model also explains higher yield of circular half-site products at early time points in concerted integration assays (37). On the other hand, concerted integration requires insertion of two viral DNA ends into target DNA (38, 39). Sequential joining of two viral DNA ends to yield the concerted integration products was noted in HIV-1 integration pathway in vitro (40). Under identical assembly conditions without DNA in solution, the RSV IN remains dimeric (**Supp Fig. 2**) suggesting that the tetramer formed in pathway II is assembled only after initial IN dimer-DNA assembly rather DNA binding to a tetrameric IN.

In conclusion, in this study, we characterized a novel intermediate in the retroviral integration pathway. These findings establish a stepwise assembly model in which RSV octameric intasome is formed through single-DNA-bound tetramers. Further studies are needed to ascertain whether an analogous pathway is present in HIV-1 and similar viruses. The approaches and assays described in this study might be useful in for dissecting assembly pathways as well as screening for novel IN inhibitors.

## Funding and additional information

This work was supported in part by the National Institutes of Health awards NIAID AI165081 and AI175583 to KKP, R35 GM149320 to EA. This work was also supported partially by Saint Louis University President Research Fund (KKP). EA would also like to acknowledge generous financial support from the Doisy Research Fund of the Edward A. Doisy Department of Biochemistry and Molecular Biology at Saint Louis University School of Medicine. The content is solely the responsibility of the authors and does not necessarily represent the official views of the National Institutes of Health.

## Conflict of interest

The authors declare that they have no conflicts of interest with the contents of this article.

## Data availability

All data needed to evaluate the conclusions in the paper are present in the paper or the supplementary materials.

## Abbreviations

IN: integrase
RSV: Rous sarcoma virus
HIV-1: human immunodeficiency virus type-1
PFV: prototype foamy virus
MMTV: mouse mammary tumor virus
MVV: maedi-visna virus
CSC: cleaved synaptic complex
STC: strand transfer complex
CIC: conserved intasome core
NTD: N-terminal domain
CCD: catalytic core domain
CTD: C-terminal domain
LTR: long terminal repeat
INSTI: integrase strand transfer inhibitor
smFRET: single-molecule FRET
TIRFm: total internal reflection fluorescence microscopy
wt: wild type
aa: amino acids
SEC: size exclusion chromatography
MW: molecular weight
AU: arbitrary units

## Notes

### Competing Interest Statement

The authors have declared no competing interest.

