## Supplementary Data for "Delineation of a novel assembly intermediate in retroviral integration pathway"

### Supp Fig. 1 Strand transfer assay

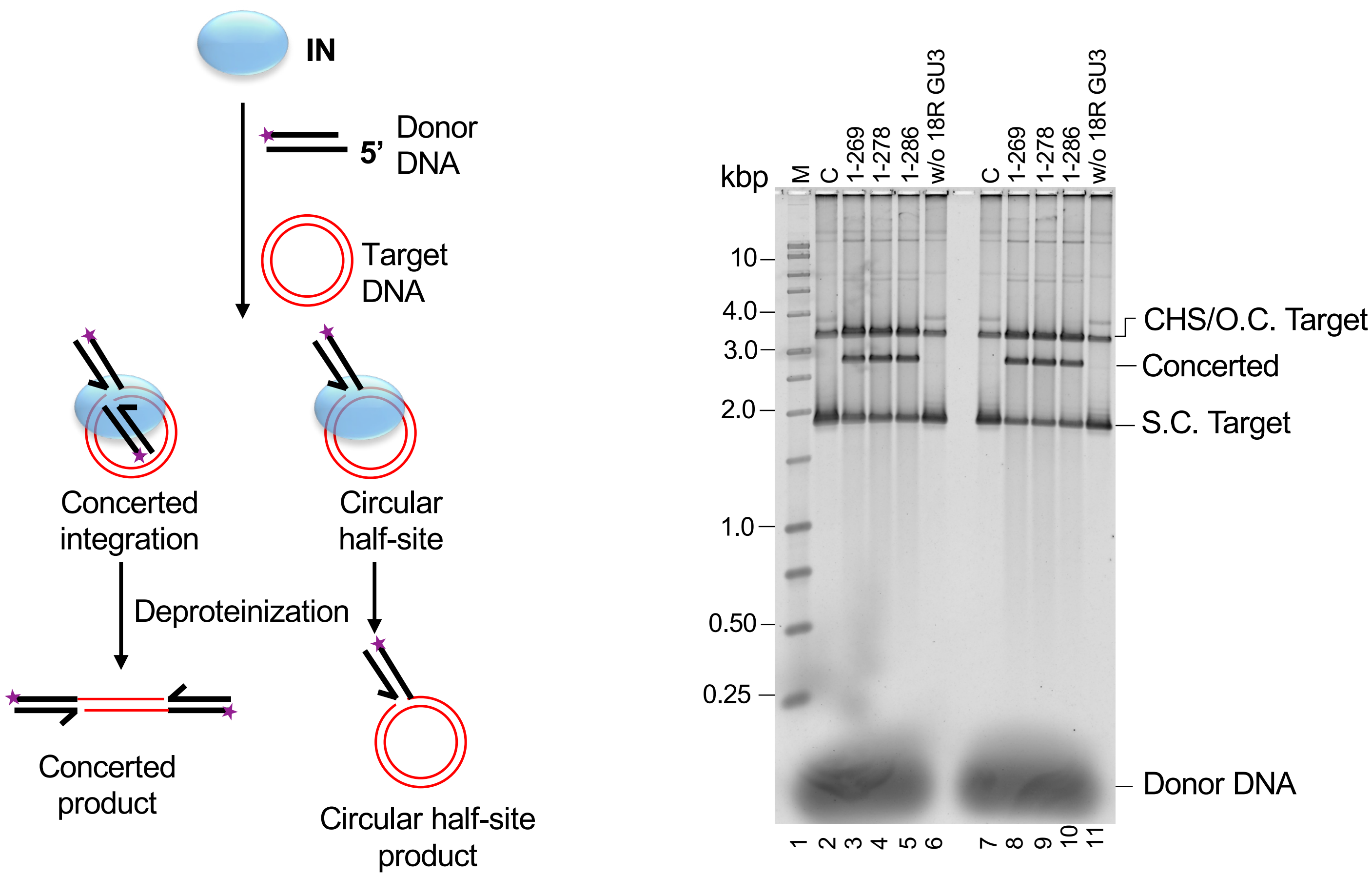


**Supp Fig. 1. Strand transfer activities of RSV IN. A.** Schematic of strand transfer assay. **B.** The concerted activity of RSV IN with different truncations of C-terminal tail region was determined with 18R GU3 donor substrate at 37°C for 15 (lanes 2-6) and 30 min (lanes 7-11). The products were deproteinized and run on a 1.3% agarose gel. Lane 1, marked M contains the molecular size marker (Promega kb ladder). Lane 2 and 7 marked C does not contain IN. lanes 3-5 and 8-10 contain different IN as indicated on the top. Lanes 6 and 11 did not contain viral DNA substrate. CHS, circular half-site; s.c. supercoiled; o.c. open circular. Unused donor DNA is indicated at bottom right.

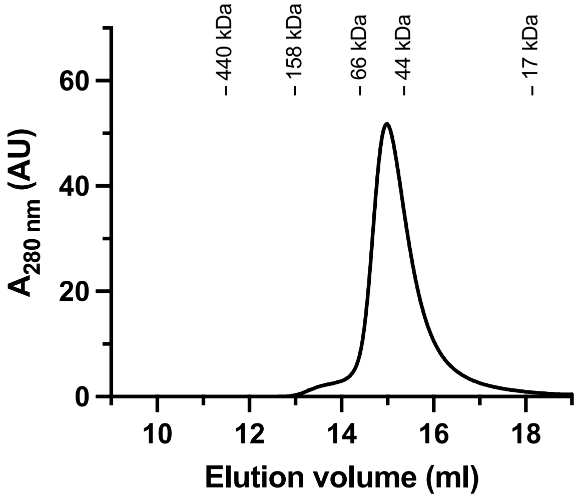


**Supp Fig. S2**. **RSV IN is predominantly dimeric.** wt RSV IN was analyzed by SEC using Superdex 200 Increase column (10 x 300 mm). The IN (45 μM) was incubated in the CSC intasome assembly buffer overnight at 18°C before SEC analysis. IN eluted predominantly as dimeric species. The molecular weight standards were run in parallel with each analysis and their elution positions are marked.

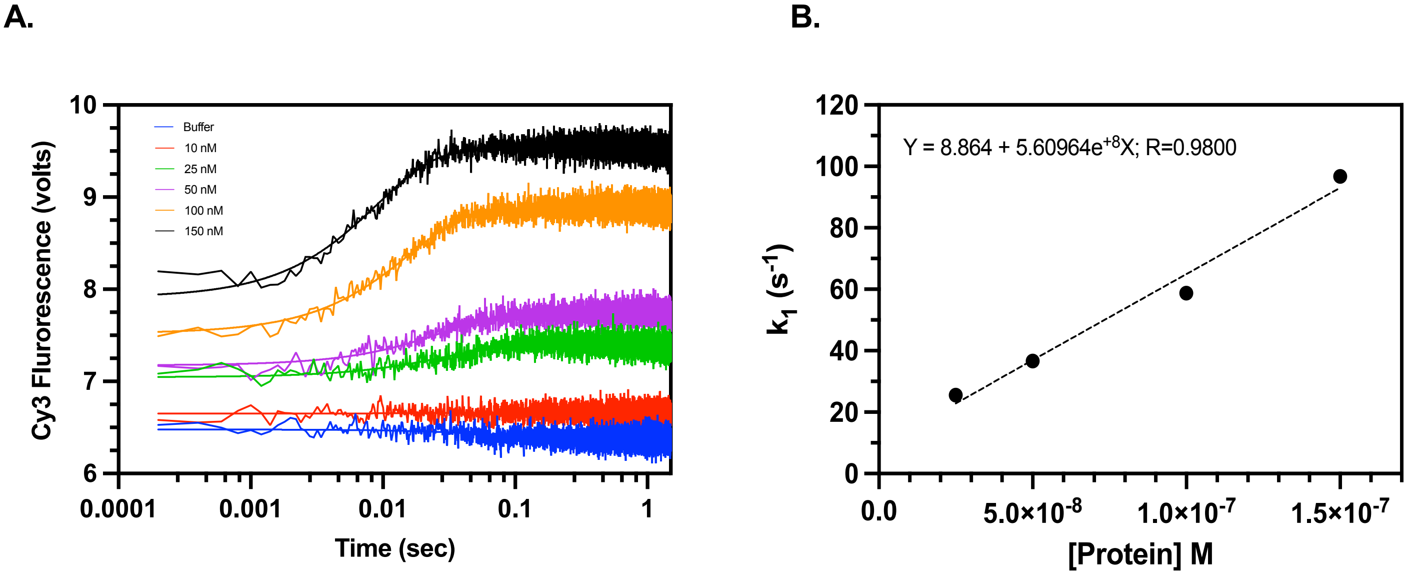


**Supp Fig. S3. Stopped flow kinetics to determine RSV IN – LTR DNA interactions. A.** Cy3 labeled 18R-GU3 (50 nM) was mixed with increasing concentrations of RSV IN and change in Cy3 fluorescence was measured. The concentrations indicated are post-mixing. **B.** Binding rate vs protein concentration fitted using simple linear regression in GraphPad Prism. The dissociation constant for IN-DNA interaction was determined to be 15.8 nM.

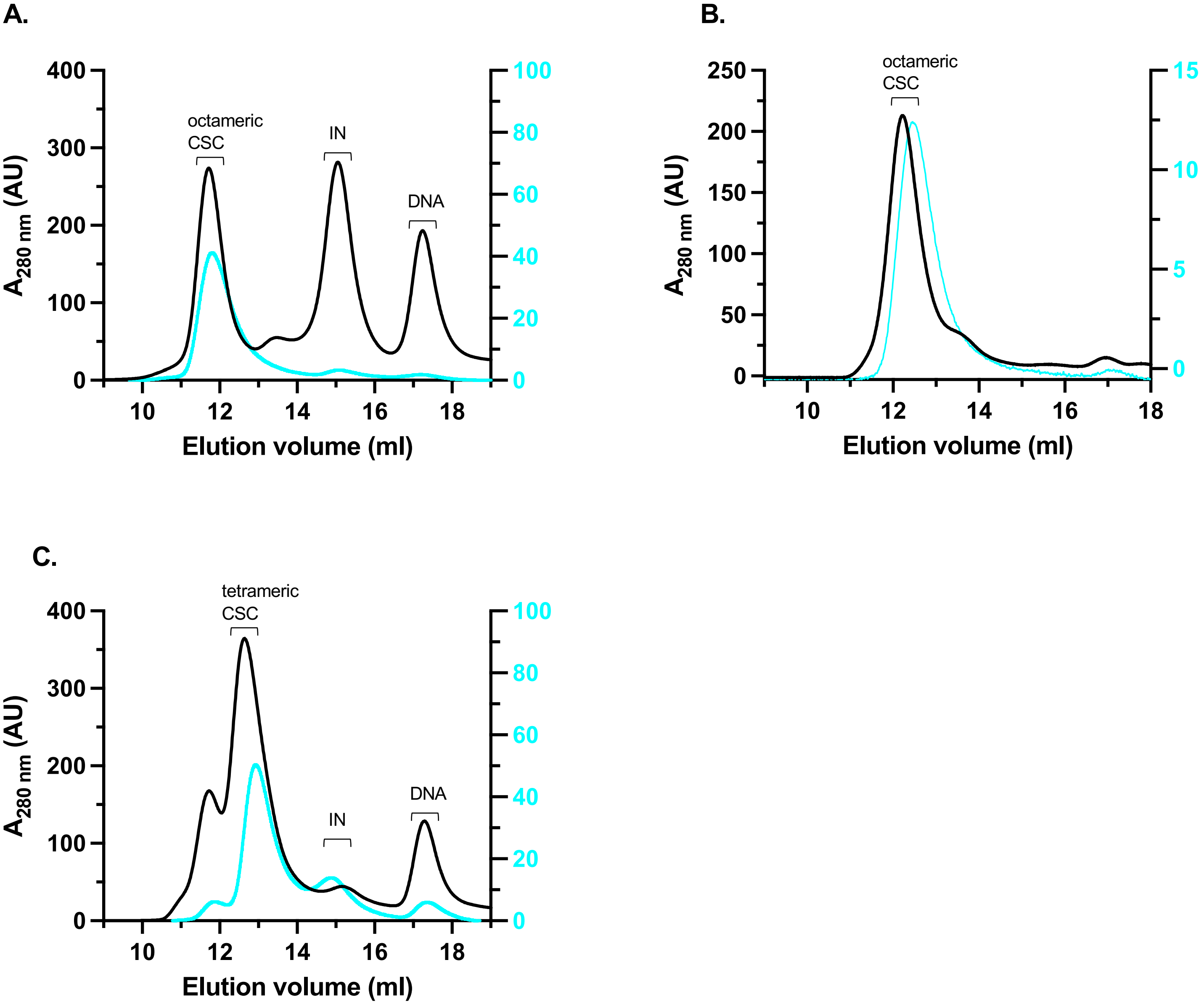


**Supp Fig. S4. Stability of RSV CSC.** Intasomes were assembled and purified by SEC (indicated by black line). Peak fractions were pooled and rechromatographed (shown in cyan) after incubation at 22°C for 30 min to determine the stability. **A.** The octameric intasome without drug. **B.** Octameric intasome formed in the presence of INSTI MK-2048. **C.** Tetrameric intasome formed in the presence of INSTI MK-2048. Right Y-axis shows the $A_{280\ nm}$ (AU) for the rechromatographed samples.
